# Alteration of menaquinone isoprenoid chain length and antibiotic sensitivity by single amino acid substitution in HepT

**DOI:** 10.1101/2020.05.11.089490

**Authors:** Suresh Panthee, Atmika Paudel, Hiroshi Hamamoto, Anne-Catrin Uhlemann, Kazuhisa Sekimizu

## Abstract

**Objectives:** *Staphylococcus aureus* Smith strain is a historical strain widely used for research purposes in animal infection models for testing the therapeutic activity of antimicrobial agents. We found that it displayed higher sensitivity towards lysocin E, a menaquinone (MK) targeting antibiotic, compared to other *S. aureus* strains. Therefore, we further explored the mechanism of this hypersensitivity.

**Methods:** MK production was analyzed by high-performance liquid chromatography and mass spectrometric analysis. *S. aureus* Smith genome sequence was completed using a hybrid assembly approach, and the MK biosynthetic genes were compared with other *S. aureus* strains. The *hepT* gene was cloned and introduced into *S. aureus* RN4220 strain using phage mediated recombination, and lysocin E sensitivity was analyzed by the measurement of minimum inhibitory concentration and colony-forming units.

**Results:** We found that Smith strain produced MKs with the length of the side chain ranging between 8 – 10, as opposed to other *S. aureus* strains that produce MKs 7 – 9. We revealed that Smith strain possessed the classical pathway for MK biosynthesis like the other *S. aureu*s. HepT, a polyprenyl diphosphate synthase involved in chain elongation of isoprenoid, in Smith strain was unique with a Q25P substitution. Introduction of *hepT* from Smith to RN4220 led to the production of MK-10 and an increased sensitivity towards lysocin E.

**Conclusions:** We found that HepT was responsible for the definition of isoprenoid chain length of MKs and antibiotic sensitivity.

## Introduction

Menaquinone (MK), found in the cytoplasmic membrane, is an essential component of the electron transport chain in Gram-positive bacteria. Apart from respiration, it plays vital roles in oxidative phosphorylation and the formation of transmembrane potential. Given the importance of MK in cellular survival, MK and its biosynthesis has been extensively studied.^1–3^ It has been shown that MK analogs inhibit the bacterial growth^4^ and several enzymes involved in MK biosynthesis such-as isoprenoid precursor;^5^ naphthoquinone;^6^ and incorporation of the isoprenoid side chain to naphthoquinone moiety^7^ can independently be targeted for antimicrobial agent discovery against Gram-positive and acid-fast microbes. Recently, we reported that lysocin E, a non-ribosomally synthesized peptide^8, 9^ produced by *Lysobacter* sp. RH2180-5, directly targets MK in the bacterial membrane exerting rapid and potent bactericidal activity.^10^

MK is a 2-methyl-1,4-naphthoquinone with an isoprenoid side chain attached at the 3-position. MK is generally referred to as MK-n, where n denotes the number of isoprenoid units between 4 and 13 attached to the naphthoquinone core. The units of isoprene in the MKs differ among different species and sometimes even within the same species.^11^ The difference in MK isoprenoid chain formed a basis of bacterial chemotaxonomic identification in pre genomic era.^12^

*Staphylococcus aureus* is a human commensal and an opportunistic pathogen responsible for a large number of hospitalization and deaths. Global spread and rise of methicillin-resistant^13,14^

and vancomycin-resistant *S. aureus* strains ^15–17^ have added the burden to health-care systems. *S. aureus* uses MKs with the length of the side chain ranging between 7 – 9, where MK-8 is the most predominant.^12^ *S. aureus* strain Smith, isolated in 1930, is widely used in the laboratory for the development of mouse infection model as it displays a high degree of virulence against mouse model.^18^ Previously, we found that it displayed a higher susceptibility towards menaquinone targeting antibiotic-lysocin E.^10^ This led to speculation that MK biosynthetic machinery in *S. aureus* Smith might be different from other *S. aureus*. In this study, we report the complete genome sequence, MK analysis of *S. aureus* Smith and the factor responsible for its hypersensitivity towards lysocin E. To the best of our knowledge, this is the first report of the identification of *S. aureus* strain producing MK-10, and the involvement of a single substitution in HepT for MK-10 production and sensitivity towards antibiotic.

## Materials and Methods

### Microorganisms and culture conditions

The bacterial strains and plasmids used in this study are summarized in **Table 1**. *S. aureus* strains were routinely grown on tryptic soy broth, and *Escherichia coli* was grown on Luria-Bertani medium. Antibiotics were supplemented to the medium as required.

**Table 1:** Bacterial strains and plasmids used in this study.

| Strain/Plasmid | Details/Source |
| --- | --- |
| *Staphylococcus aureus* |  |
| Smith | Isolated in 1930,<sup>18</sup> obtained from ATCC13709 |
| RN4220 | Restriction deficient strain, laboratory stock |
| Newman | Isolated in 1952,<sup>19</sup> laboratory stock |
| NCTC8325-4 | Parent strain of RN4220, laboratory stock |
| JE2 | USA300 strain obtained from BEI Resources |
| MRSA4 | Clinical isolate<sup>20, 21</sup> |
| 71101 | Clinical isolate<sup>22</sup> |
| NCTC5663 | Public Health England |
| *Escherichia coli* HST08 | Competent cells for routine cloning from Takara |
| pND50-pfbaA | pND50 with *fbaA* promoter inserted in EcoRI/BamHI site<sup>23</sup> |
| pND50-pfbaA-hepT<sub>Smith</sub> | hepT<sub>Smith</sub> in pND50-pfbaA |
| pND50-pfbaA-hepT<sub>RN4220</sub> | hepT<sub>RN4220</sub> in pND50-pfbaA |

### Whole-genome sequencing, assembly and comparative genomic analysis

The complete genome of *S. aureus* Smith was sequenced using hybrid genome assembly as explained previously^24–26^ using 1 μg and 100 ng of genomic DNA for Oxford Nanopore MinION and ThermoFisher Ion PGM, respectively. The assembled genome was annotated using the NCBI Prokaryotic Genome Annotation Pipeline. The draft genome of *S. aureus* 71101 was obtained by Illumina sequencing.^22^ The complete genome sequences of 324 *S. aureus* strains were obtained from NCBI GenBank, and amino acid sequences of MK biosynthetic genes were obtained using BLAST search.

### *hepT* cloning and heterologous expression

The *hepT* gene from *S. aureus* was amplified using the primer sets BamF vs SalR1 and BamF vs SalR2 for Smith and RN4220 strains, respectively (**Table 2**). The BamHI SalI digested PCR product was then ligated to pND50-p*fbaA* vector^23^ digested with the same enzymes to construct pND50-p*fbaA-hepT*_Smith_ and pND50-p*fbaA-hepT*_RN4220_, respectively. The ligated plasmid was then transformed to *Escherichia coli* HST08 (Takara Bio) and selected on chloramphenicol plates. The strains with correct sequences were selected for transformation into electrocompetent *S. aureus* RN4220. Insertion in the RN4220 strain was then confirmed by PCR.

**Table 2:**
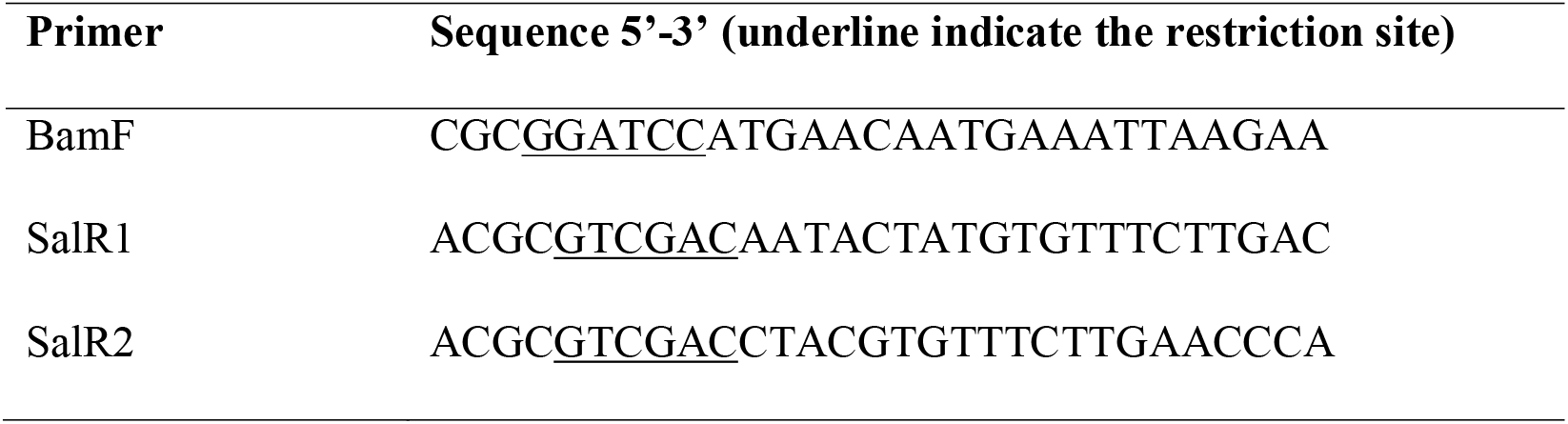
Primers used to amplify *hepT* gene.

### Menaquinone extraction and HPLC analysis

*S. aureus* strains were cultured overnight in 5 mL TSB supplemented with antibiotics as required in a shaking incubator maintained at 37°C. The full growth was then diluted 100-fold in the 5 mL TSB medium without antibiotics and incubated in the same shaker for 16 hours. A 300 μL of the culture broth was extracted twice with 1.5 mL of hexane 5: ethanol 2. The supernatant was pooled, dried *in-vacuo*, dissolved in 200 μL ethanol and 80 μL of it was analyzed using a Waters Alliance high-performance liquid chromatography (HPLC) system equipped with a Senshu Pak PEGASIL ODS SP100 column (4.6φ × 250 mm) maintained at 40°C. After the application of the sample to the column equilibrated with 1 mL◻min^−1^ of 20% diisopropyl ether in methanol, the column was eluted with the same solvent. Detection was made using a fluorescent detector using wavelengths 320 and 430 nm for excitation and emission, respectively, after post-column reduction using a platinum column.

### High resolution mass spectrometric analysis

High resolution mass spectrometric analysis was performed on a UPLC/MS system using a Waters Acquity UPLC consisting of 2.1 × 50 mm Acquity UPLC® BEH C18 1.7 μm column. After the injection of the sample to the column equilibrated with 0.3 mL◻min^−1^ of 100% methanol, the eluate was continuously applied to a Waters Xevo G2-XS QTof mass spectrometer. The data at the mass range of 100 – 1700 Da were collected in ESI positive mode using a source capillary voltage of 2.00 kV. The data were obtained using MassLynx 4.1 (Waters Milford, MA, USA) and analyzed by UNIFI Scientific Information System (Waters).

### Lysocin E susceptibility

Clinical and Laboratory Standards Institute broth microdilution method was used to determine the minimum inhibitory concentrations (MIC).^27^ Briefly, serial dilutions of lysocin E were prepared in cation-adjusted Mueller-Hinton Broth (Difco, Franklin Lakes, NJ, USA) and a 100 μL aliquot was then dispensed to each well of a 96-well plate. Inoculum containing approximately 1×10^6^ colony forming units (CFU)/mL of bacteria was prepared from *Staphylococci* colonies grown at 37°C on Tryptic Soy Broth (Difco) agar plates. 10 μL of it was added to each well of the 96-well plate and incubated at 37°C for 18 h. The minimum concentration that inhibited the growth of bacteria was considered as the MIC value.

Viability of *S. aureus* upon treatment with lysocin E was determined as described previously ^21, 28^ following NCCLS protocol.^29^ Briefly, the overnight full growth of *Staphylococci* was diluted 100 fold with 5 mL TSB and incubated at 37°C with shaking. After the OD_600_ reached 0.1, 1 mL aliquot was collected and treated with 1 mg/L of lysocin E, and incubation was continued for 30 minutes. The number of the surviving bacteria was counted by spreading on Mueller Hinton agar plates. Untreated samples at time zero were considered as 100% and used to calculate percentage survival.

## Results and Discussion

### Higher Sensitivity of *S. aureus* Smith towards lysocin E

Lysocin E (**Figure 1a**) is a recently discovered antibiotic effective against Gram-positive bacteria that utilize MK for respiration.^10, 30^ Lysocin E has a potent and rapid bactericidal activity. It has a minimum inhibitory concentration (MIC) value of 4 mg/L against most of the laboratory *S. aureus* strains, which we tested, except for Smith strain, against which lysocin E consistently displayed an MIC value of 2 mg/L (**Figure 1b**). We further found a more potent bactericidal activity of lysocin E against Smith compared to Newman and JE2 strains (**Figure 1c**), suggesting its hypersensitive nature. As lysocin E targets MK,^10^ and *S. aureus* has MK as the sole quinone known to be utilized for respiration,^31^ we speculated that the MKs in Smith strain could be different from other *S. aureus* strains. However, there is no study about the type, content, and biosynthesis of MKs in *S. aureus* Smith. Therefore, we extracted MKs form the overnight cultures of the *S. aureus* Smith, Newman, and JE2 strains and analyzed by HPLC. Consistent with the previous report,^32^ Newman strain mainly produced MK-7 and MK-8, MK-8 being the most abundant, and trace amounts of MK-9. While MK production in JE2 was similar to that of Newman strain, Smith strain mainly produced MK-8 and MK-9, with MK-9 being the most abundant, and there appeared an undefined peak at the retention time of 34 minutes (**Figure 2a, b**). We then extracted MKs from a 50-mL volume of culture and separately collected each peak and analyzed by high-resolution mass spectrometry. We found that the peaks were 739.5449, 807.6043 and 875.6648 corresponding with [M+Na]^+^ of MK-8, MK-9, and MK-10, respectively (**Figure 2c**). The undefined peak was thus identified as MK-10. Therefore, as opposed to the major quinone MK-8 in *S. aureus,*^12^ Smith strain produced MK-9 predominantly. In addition, Smith strain produced MK-10, an MK that has not been reported in *S. aureus*. These results suggested that longer chain MKs in Smith strain might be responsible for its hypersensitivity towards lysocin E. Previously we found that *S. aureus* strains with mutation and/or deletions in the genes involved in MK biosynthesis were resistant to lysocin E^10^ suggesting that analysis of MK biosynthetic genes in Smith would give an insight upon its hypersensitivity.

**Figure 1:**
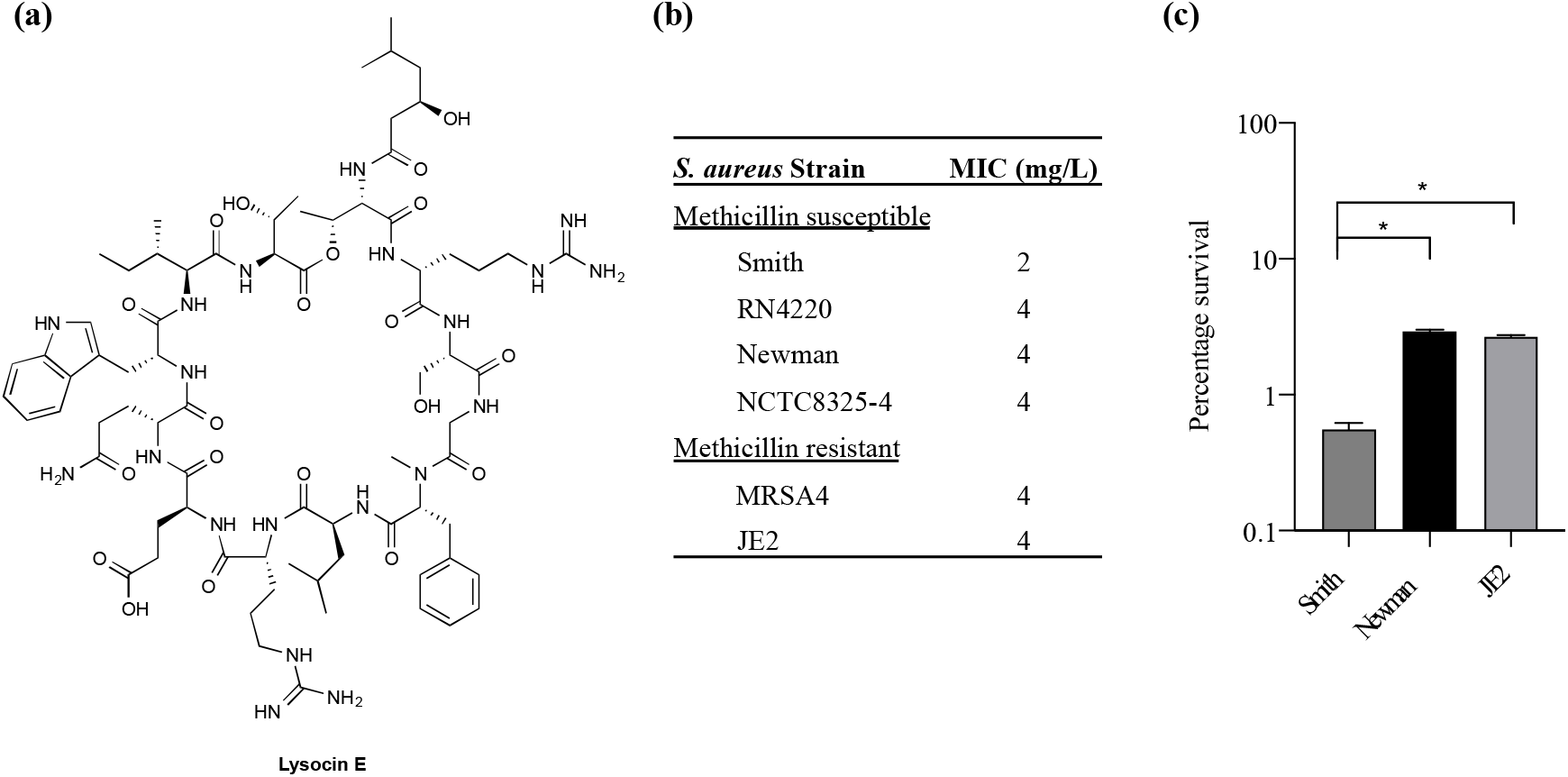
Lysocin E and its antimicrobial activity. (a) Chemical structure of lysocin E. (b) Minimum inhibitory concentrations of lysocin E against various *S. aureus*. MIC was determined by broth microdilution assay and represented as the median value obtained from 10 experiments. **(c)** Bactericidal activity of lysocin E.*S. aureus* strains were treated with 1 mg/L lysocin E for 30 minutes, and bacterial viability was determined. Triplicate data are represented as mean ± SEM and statistical analysis was performed by one-way ANOVA using Dunnett’s multiple comparison test in GraphPad Prism. The asterisk indicates a *p-value* of <0.0001.

**Figure 2:**
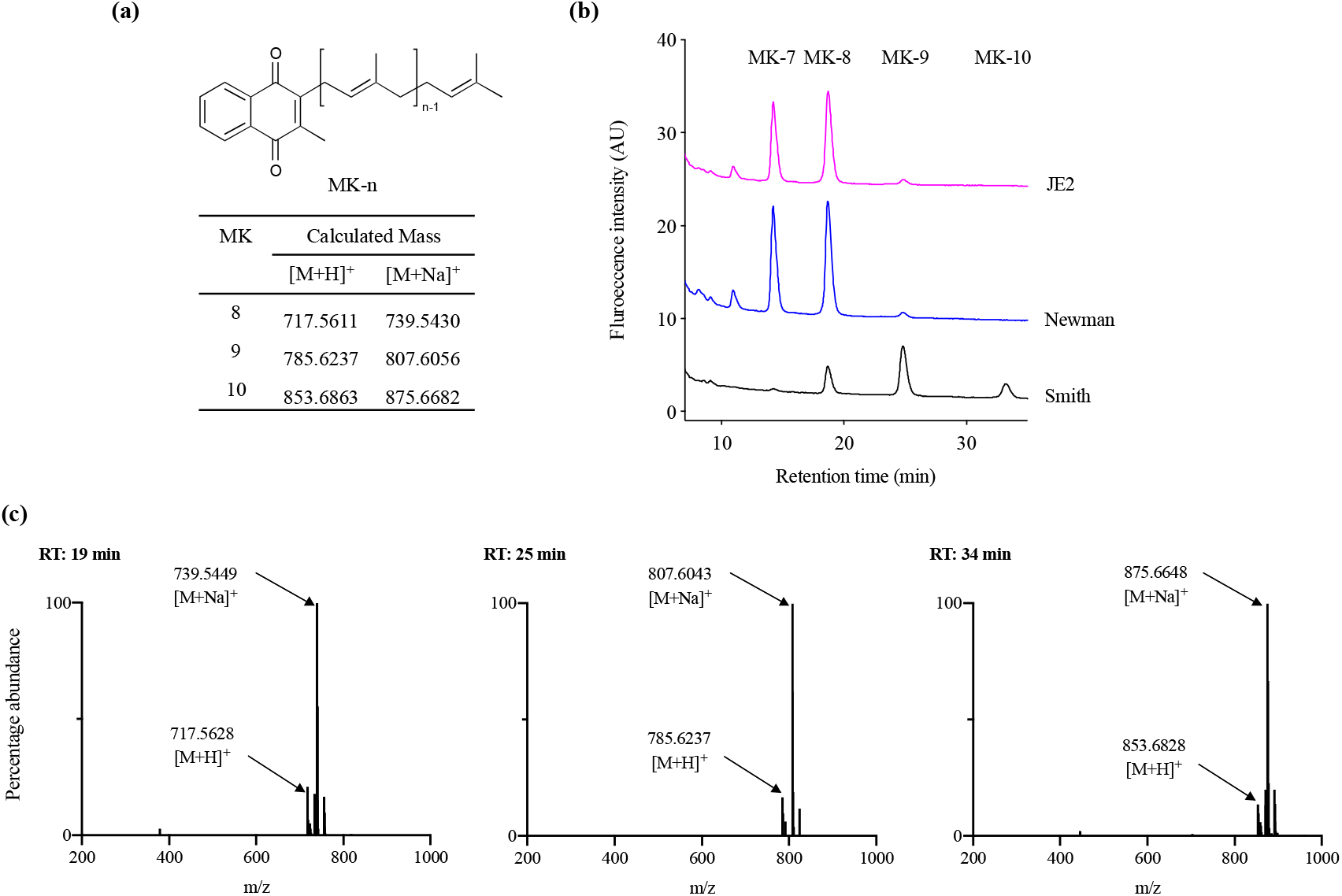
Analysis of MKs from *S. aureus*. **(a)** Chemical structure of MK-n and calculated exact mass of MK-8, 9, and 10 in positive ion analysis. **(b)** Analysis of MK extract from *S. aureus* Smith, Newman, and JE2. **(c)** High-resolution mass spectrometric analysis of peaks that appeared in Smith at 19, 25 and 34 minutes.

### Analysis of MK biosynthetic pathway in *S. aureus* Smith

The ability of the Smith strain to produce MK-10 and an association of mutations in MK biosynthetic genes with lysocin E resistance^10^ triggered us to analyze the MK biosynthetic pathway of this strain so that we could identify the genetic basis of this unique feature. We obtained the complete genome sequence of the Smith strain using a hybrid Ion PGM and Nanopore MinION sequencing approach^24, 26^. We performed a BLAST search against the genes involved in MK biosynthetic pathway. We found that the Smith strain harbored orthologs of all the genes involved in the classical pathway (**Figure 3**). We further aligned 11 MK biosynthetic enzymes among Newman, JE2 and Smith strains to find that Newman and JE2 shared an end to end sequence identity in all the enzymes, while Smith strain had amino acid substitution(s) in enzymes except MenA, MenG, and MenI (**Figure 3, Supplementary Figure S1**).

**Figure 3:**
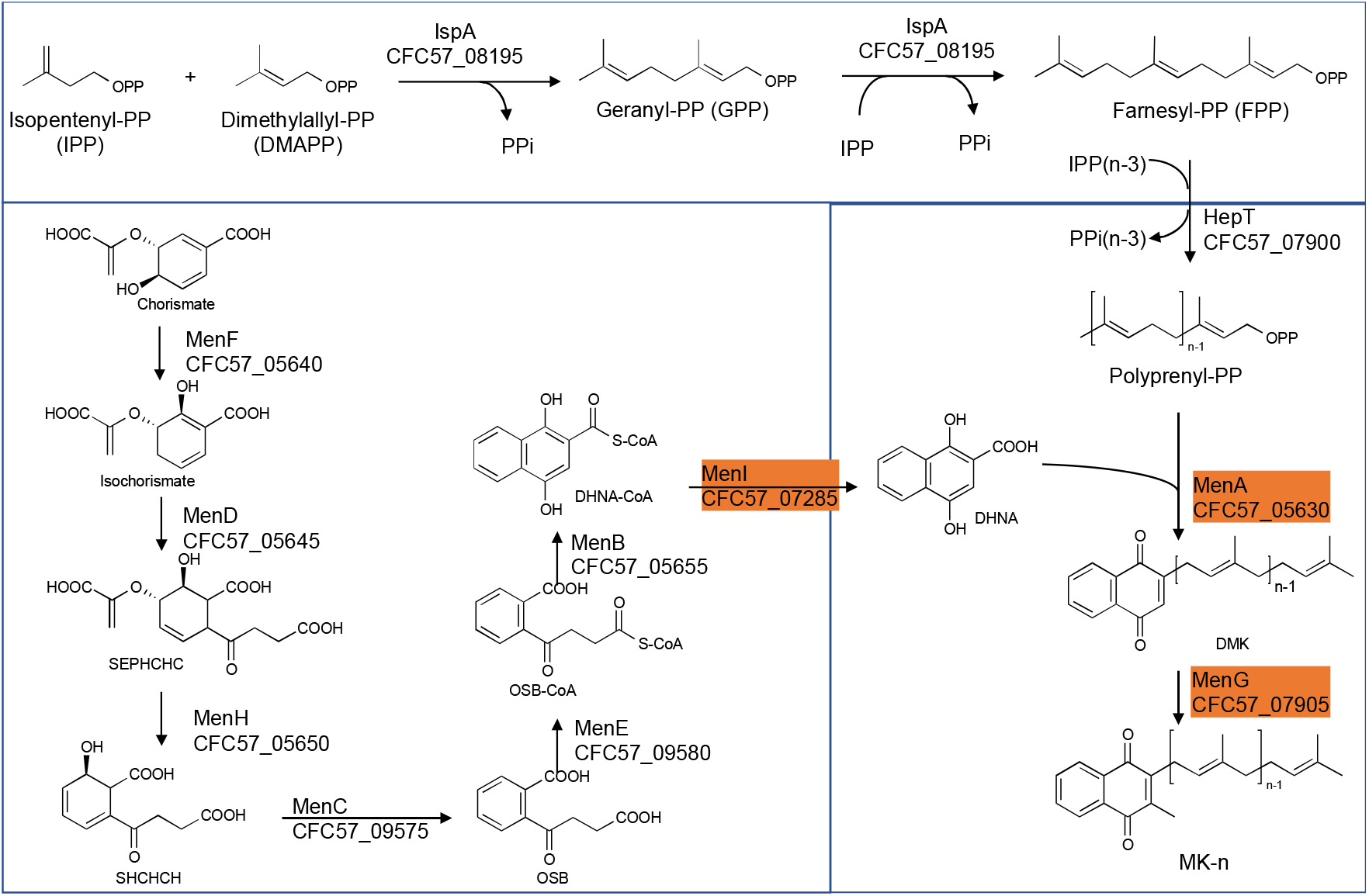
The classical MK biosynthetic pathway in *S. aureus* Smith. The highlighted enzymes have an end to end sequence identity between *S. aureus* Smith, JE2 and Newman strains.

Among the Smith MK biosynthetic enzymes that harbored amino acid substitution(s), the majority were involved in the formation of 1,4-dihydroxy-2-naphthoate. Among the enzymes involved in isoprenoid side chain biosynthesis, IspA (CFC57_08195) and HepT (CFC57_07900) had 2, and 3, amino acid substitutions, respectively. IspA is predicted to be involved in the formation of Farnesyl-PP, and HepT is predicted to be involved in the condensation of Isopentenyl-PPs and Farnesyl-PPs, resulting in the formation of all-trans-polyprenyl-PP. Based on this, we speculated that Smith HepT (HepT_Smith_ now onwards) might be involved in the formation of longer chain polyprenyl-PPs to be attached to 1,4-dihydroxy-2-naphthoate by MenA (CFC57_05630).

### Analysis of Staphylococcal HepT involved in polyprenyl diphosphate biosynthesis

We then analyzed the HepT sequence of all *S. aureus* strains whose complete genome sequence was available in NCBI. We focused on three substitutions (Pro-25, Leu-170, and Asp-288) that were different in Smith strain from Newman and JE2 strains (**Figure 4a**) and found that the HepT from 325 *S. aureus* strains could be categorized to five types which we named type 1 to type 5. Type 1 – 4 were present in at least 10 strains while type 5 was unique for Smith strain with Pro-25 (**Figure 4b**). Among these, we analyzed the MK content from strains harboring four available types of HepT and found that only Smith could produce MK-10 (**Figure 4c**). This result suggests that Pro-25 of HepT_Smith_ could be responsible for longer chain MK biosynthesis.

**Figure 4.**
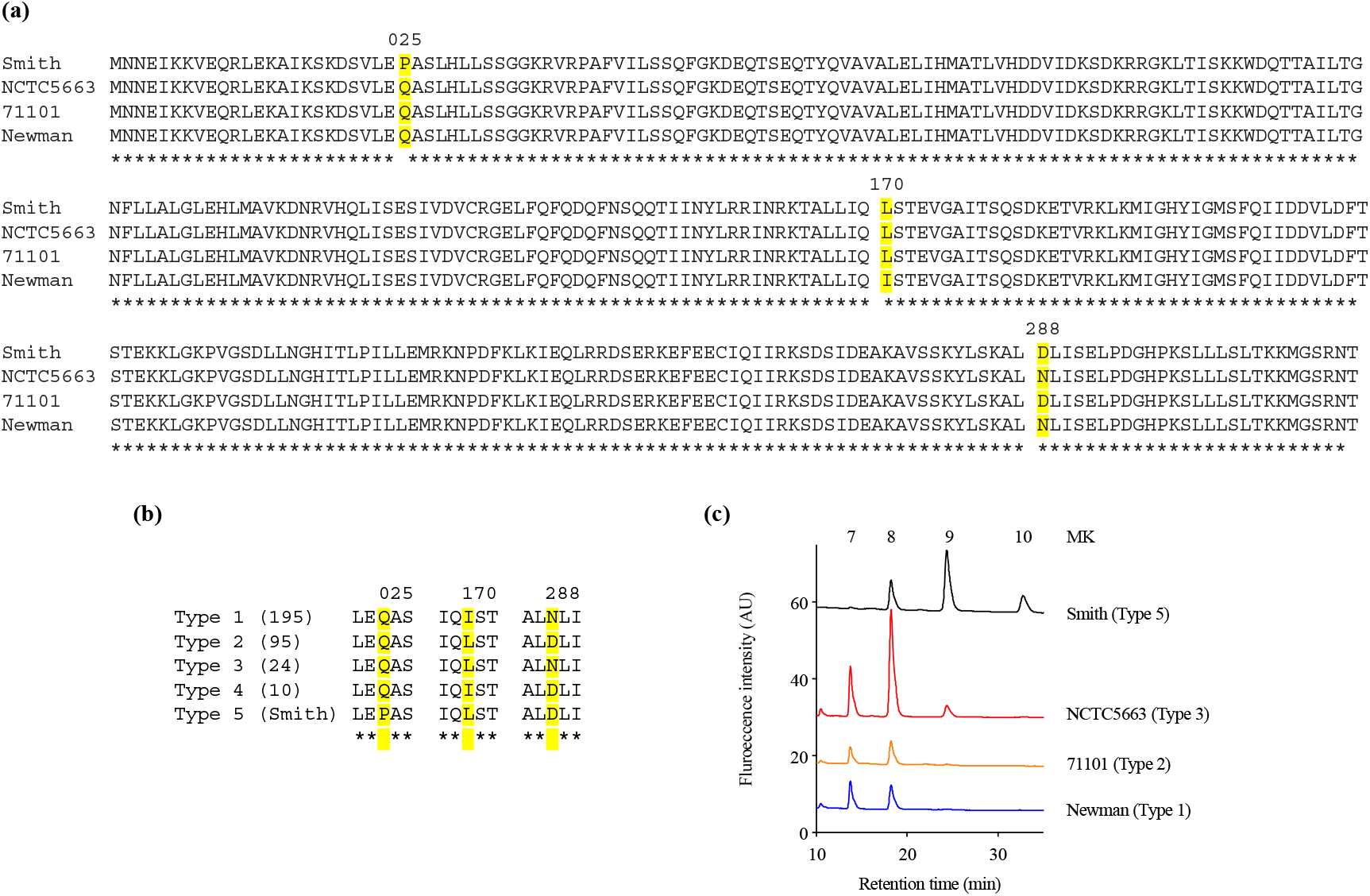
Analysis of Staphylococcal HepTs. **(a)** Alignment of HepT from strains Smith, NCTC5663, 71101, and Newman. **(b)** Five types of *S. aureus* based on the position of amino acids at 25, 170, and 288 in the HepT sequence. Numbers in parenthesis indicate the number of strains in each type. Type 5 only contained Smith strain. **(c)** MK content of representative *S. aureus* strains to harbor four HepT types.

### HepT_Smith_ is involved in chain length determination of MK

To confirm the role of HepT_Smith_ in longer chain MK biosynthesis, we cloned the *hepT* gene from the Smith strain and expressed it under the control of the constitutive expression promoter.^23^ The plasmid thus obtained was introduced into the restriction deficient strain *S. aureus* RN4220. We also cloned the *hepT* gene from the RN4220 strain and introduced it into the RN4220 strain. We compared the MK production among Smith strain, RN4220 with empty vector, *hepT*_Smith_, and *hepT*_RN4220_ While the production of shorter chain MKs (MK-7 and MK-8) were similar in all the transductants, the introduction of *hepT*_Smith_ in RN4220 resulted in significantly higher production of MK-9 and the appearance of MK-10 (**Figure 5a-c**). RN4220 harboring empty vector or *hepT*_RN4220_ predominantly produced MK-7 and MK-8, with a trace amount of MK-9, and the MK pattern was indifferent from that of the wild type strain (**Figure 5a-c**). These results suggest that HepT_Smith_ is responsible for the biosynthesis of longer chain MKs.

**Figure 5.**
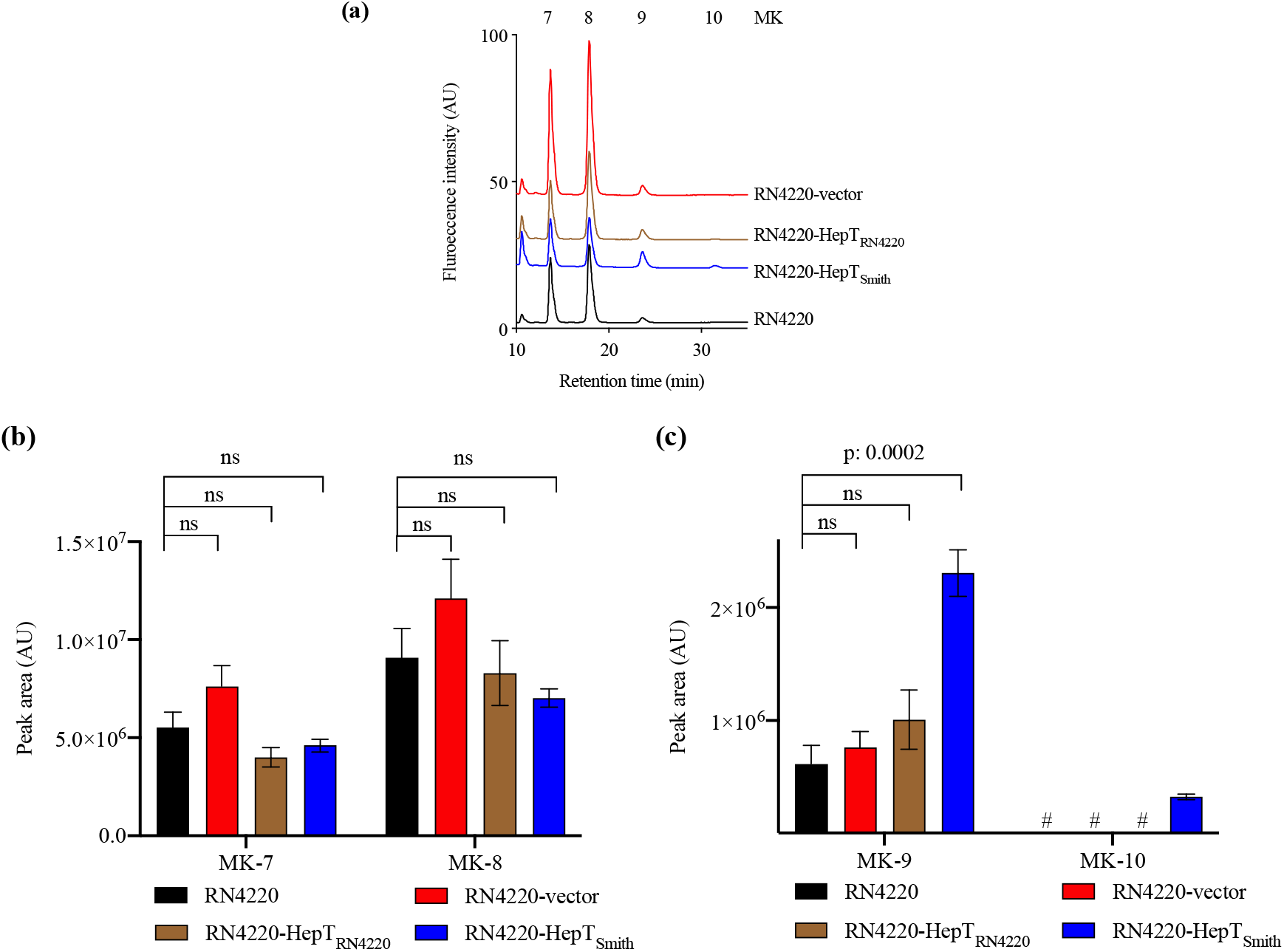
Analysis of MKs from *S. aureus* RN4220 with heterologously expressed HepT. **(a)** Representative HPLC chromatograms. **(b)** Peak area of MK-7 and MK-8. **(c)** Peak area of MK-9 and MK-10. Data are from three independent experiments and represented as mean ± SEM. Statistical analysis was performed by one-way ANOVA using Dunnett’s multiple comparison test in GraphPad Prism, and a *p-value* less than 0.05 was considered significant. ns: non-significant. # indicates an undetectable amount of MK-10.

### Longer chain MKs are responsible for hypersensitivity to lysocin E

The hypersensitivity of Smith strain towards lysocin E, the presence of MK-10 in Smith strain, its unique HepT, and evidence showing the involvement of HepT_Smith_ in MK-10 production led us further to explore the role of HepT_Smith_ in lysocin E sensitivity. To elucidate this, we compared the viability of Smith and RN4220 strains harboring the empty vector and HepT_Smith_ upon treatment with 1 mg/L of lysocin E to find that a 30 minutes treatment drastically reduced the number of viable bacteria (**Figure 6**). Furthermore, Smith and RN4220 expressing HepT_Smith_ were hypersensitive to lysocin E treatment, suggesting that increased production of MKs harboring longer isoprenoid side chain might be responsible for the phenomena.

**Figure 6.**
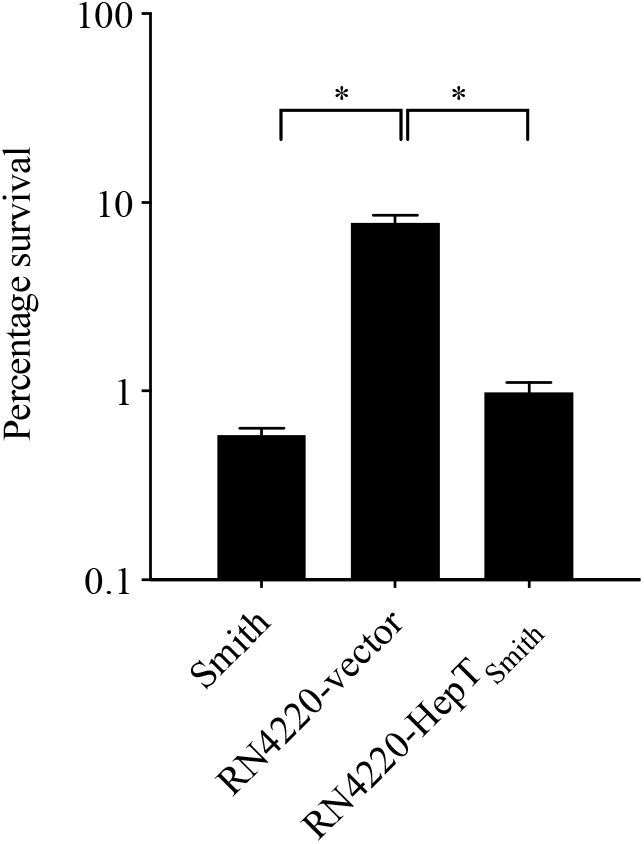
Survival of *S. aureus* in the presence of lysocin E. Exponentially growing bacteria were treated with 1 mg/L of lysocin E for 30 min, and the colony-forming units were counted Triplicate data are represented as mean ± SEM. Statistical analysis was performed by one-way ANOVA using Dunnett’s multiple comparison test in GraphPad Prism, and the asterisk indicates a *p*-value of <0.0001.

In addition to MK biosynthesis, isoprenoids are critical for the biosynthesis of membrane lipids, carotenoids, sterols, and other components of the bacterial cell wall.^33^ Isopentenyl-PP, one of the substrates of HepT and the starting molecule for other isoprenoid biosynthesis, is synthesized either via 2-C-methyl-D-erythritol-4-phosphate (MEP) and/or mevalonate pathway.^34, 35^ The enzymes of the MEP pathway have been used as targets for antibiotic discovery against microbes that harbor the MEP pathway.^5, 36^ Given that *S. aureus* relies on the mevalonate pathway,^37^ HepT or other enzymes from this pathway can be targeted for the antistaphylococcal drug development.^38, 39^

In summary, we completed the genome sequence of *S. aureus* Smith and performed the genomic analysis of the MK biosynthetic pathway to show that a classical pathway for MK biosynthesis is present in this strain. We demonstrated that Pro-25 substitution in HepT was responsible for longer chain MK biosynthesis, and this was associated with hypersensitivity towards lysocin E. This indicated that lysocin E might disrupt the bacterial membranes containing longer chain MKs more efficiently which requires further analysis. To the best of our knowledge, this is the first report of the identification of *S. aureus* strain producing MK-10.

## Acknowledgements

The JE2 strain was provided by the Network on Antimicrobial Resistance in *Staphylococcus aureus*, distributed by BEI Resources, National Institute of Allergy and Infectious Diseases, National Institutes of Health. We thank Ms. Maeda from Genome Pharmaceuticals Institute Co., Ltd. for technical assistance.

## Transparency declarations

None to declare.

## Supplementary data

Supplementary figure 1 is available online.

## Data Availability

The complete genome assembly of *S. aureus* Smith has been deposited at DDBJ/ENA/GenBank with accession numbers: CP029751 and CP029750, for chromosome and plasmid pSS41, respectively. The BioProject accession number for this project is PRJNA392199.

