## Supplementary Figure S1 for "Alteration of menaquinone isoprenoid chain length and antibiotic sensitivity by single amino acid substitution in HepT"

**Running Title: HepT in MK biosynthesis and antibiotic sensitivity**

Suresh Panthee[1,§], Atmika Paudel[1,§], Hiroshi Hamamoto[1], Anne-Catrin Uhlemann[2], and Kazuhisa Sekimizu[1,*]

[1] Teikyo University Institute of Medical Mycology, Otsuka 359, Hachioji, Tokyo 192-0395, Japan

[2] Department of Medicine, Columbia University Medical Center, New York, New York, USA

§ These authors contributed equally to this work.

### Supplementary information

```
IspA     198      270
Smith  DDATK    ELMQID
Newman DDTTK    ELTQID
JE2    DDTTK    ELTQID
       **:**    ** ***

HepT     025      170      288
Smith  LEPAS    IQLST    ALDLI
Newman LEQAS    IQIST    ALNLI
JE2    LEQAS    IQIST    ALNLI
       ** **    **:**    **:**

MenF     273
Smith  DKMPK
Newman DKTPK
JE2    DKTPK
       ** **

MenD     274      537
Smith  DLKVD    RENNF
Newman DLNVD    REDNF
JE2    DLNVD    REDNF
       **:**    **:**

MenH   1                                         43   180    221
Smith  MTHYNYYEANVETNQVLVLLHGFLSDSRTYHNHIDKYTDICHV   SPQKM  FVKIA
Newman MTHYKFYEANVETNQVLVFLHGFLSDSRTYHNHIEKFTDNYHV   SPHKM  FVQIA
JE2    MTHYKFYEANVETNQVLVFLHGFLSDSRTYHNHIEKFTDNYHV   SPHKM  FVQIA
       ****::************:**************:*:**   **   **:**  **:**

MenC     013      074    189                   211    265    306  311
Smith  YSVPF    FEYNR    REHILYIEEPFKDISMLDEVVDG    HGVKV    VVTKSG
Newman YSEPF    FEDNR    REQVLYIEEPFKDISMLDEVADG    HGAKV    VVAHSG
JE2    YSEPF    FEDNR    REQVLYIEEPFKDISMLDEVADG    HGAKV    VVAHSG
       ** **    ** **    **::***************.**    **.**    **::**

MenE     043     090    123                       148   196     273    359     422    447
Smith  LKDYQ   QMRSI    ITMNGLLDNTMDIQFDTSNETVVSKE   KENLG   PYDLQ  MNGYL   GIIDA  ESNIS
Newman LKAYQ   QMKSI    ITTNSLLDNTMGIQYETSNETVVPKE   KESLG   PYNLQ  MNVYL   GISDA  ESDIS
JE2    LKAYQ   QMKSI    ITTNSLLDNTMGIQYETSNETVVPKE   KESLG   PYNLQ  MNVYL   GISDA  ESDIS
       ** **   **:**    ** *.******.**::******* **   **.**   **:**  ** **   ** **  **:**

MenB     196
Smith  PLDKV
Newman PLEKV
JE2    PLEKV
       **:**
```

**Supplementary Figure S1:** Alignment of MK biosynthetic genes among three *S. aureus* strains: Smith, JE2 and Newman. The conserved sequence is highlighted.
